# Vht hydrogenase is required for hydrogen cycling during nitrogen fixation by the non-hydrogenotrophic methanogen *Methanosarcina acetivorans*

**DOI:** 10.1101/2021.10.12.464174

**Authors:** Jadelyn M. Hoerr, Ahmed E. Dhamad, Thomas M. Deere, Melissa Chanderban, Daniel J. Lessner

## Abstract

*Methanosarcina acetivorans* is the primary model to understand the physiology of methanogens that do not use hydrogenase to consume or produce hydrogen (H_2_) during methanogenesis. The genome of *M. acetivorans* encodes putative methanophenazine-reducing hydrogenases (Vht and Vhx), F_420_-reducing hydrogenase (Frh), and hydrogenase maturation machinery (Hyp), yet cells lack significant hydrogenase activity under all growth conditions tested to date. Thus, the importance of hydrogenase to the physiology of *M. acetivorans* has remained a mystery. *M. acetivorans* can fix dinitrogen (N_2_) using nitrogenase that is documented in bacteria to produce H_2_ during the reduction of N_2_ to ammonia. Therefore, we hypothesized that *M. acetivorans* uses hydrogenase to recycle H_2_ produced by nitrogenase during N_2_ fixation. Results demonstrate that hydrogenase expression and activity is higher in N_2_-grown cells compared to cells grown with fixed nitrogen (NH_4_Cl). To test the importance of each hydrogenase and the maturation machinery, the CRISPRi-dCas9 system was used to generate separate *M. acetivorans* strains where transcription of the *vht, frh, vhx*, or *hyp* operons is repressed. Repression of *vhx* and *frh* does not alter growth with either NH_4_Cl or N_2_ and has no effect on H_2_ metabolism. However, repression of *vht* or *hyp* results in impaired growth with N_2_ but not NH_4_Cl. Importantly, H_2_ produced endogenously by nitrogenase is detected in the headspace of culture tubes containing the *vht* or *hyp* repression strains. Overall, the results reveal that Vht hydrogenase recycles H_2_ produced by nitrogenase that is required for optimal growth of *M. acetivorans* during N_2_ fixation.

**IMPORTANCE:** The metabolism of *M. acetivorans* and closely related Methanosarcinales is thought to not involve H_2_. Here we show for the first time *M. acetivorans* is capable of H_2_ cycling like hydrogenotrophic Methanosarcinales (e.g., *Methanosarcina barkeri*). However, unlike *M. barkeri* hydrogenase activity and H_2_ cycling is tightly regulated in *M. acetivorans* and is only utilized during N_2_ fixation to consume H_2_ production endogenously by nitrogenase. The *in vivo* production of H_2_ by nitrogenase during N_2_ reduction is also demonstrated for the first time in a methanogen. Overall, the results provide new insight into the evolution and diversity of methanogen metabolism and new details about methanogen nitrogenase that could be leveraged for practical applications, such as nitrogenase-dependent production of H_2_ as a biofuel.

## INTRODUCTION

Methanogenesis by anaerobic archaea (methanogens) is the primary source of biologically produced methane (CH_4_) and is the last step in metabolism of biomass in many anaerobic environments. Most methanogens produce CH_4_ from the reduction of carbon dioxide (CO_2_) using hydrogen gas (H_2_). Methanogens that lack cytochromes (e.g., *Methanococcus* sp.) are primarily restricted to the CO_2_-reduction pathway of methanogenesis. However, it is estimated that only one-third of CH_4_ produced by methanogens is from the reduction of CO_2_, while two-thirds is derived from the methyl group of acetate [1]. Only two genera of cytochrome-containing methanogens, *Methanosarcina* and *Methanothrix*, can produce CH_4_ with acetate [2, 3]. *Methanosarcina* sp. are the most metabolically diverse, capable of using methylated substrates, including methanol, methylamines, and methylsulfides, in addition to acetate and H_2_/CO_2_ [1]. The pathways for methylotrophic and aceticlastic methanogenesis have been extensively studied in freshwater isolates *Methanosarcina barkeri* and *Methanosarcina mazei* and the marine isolate *Methanosarcina acetivorans*. Although the overall pathways are similar, the electron transfer and energy conservation steps are different between the species [4]. For example, methanogenesis by *M. barkeri* with all substrates is H_2_ dependent, whereas methanogenesis by *M. acetivorans* with all substrates is H_2_ independent.

During methylotrophic methanogenesis (the pathway used in this study), both *M. barkeri* and *M. acetivorans* oxidize a methyl group to CO_2_ by a reversal of the CO_2_ reduction pathway and use the six electrons to indirectly reduce three additional methyl groups to CH_4_ (**Figs. 1 and 2**) [4, 5]. Both respiratory pathways involved in energy conservation require the reduction of methyl-coenzyme M (CH_4_-SCoM) to CH_4_ using coenzyme B (HSCoB) as the direct electron donor, resulting in the formation of a heterodisulfide of both coenzymes (CoMS-SCoB). In both species, CoMS-SCoB serves as the terminal electron acceptor and is reduced by a membrane-bound electron transport chain consisting of the membrane-soluble electron carrier methanophenazine and the cytochrome-containing heterodisulfide reductase (HdrDE) complex. The electron source and enzyme used to reduce methanophenazine is different between the two species. *M. barkeri* primarily utilizes a H_2_:CoMS-SCoB oxidoreductase system where H_2_ serves as an intermediate electron carrier (**Fig. 1**), while *M. acetivorans* primarily utilizes a F_420_H_2_:CoMS-SCoB oxidoreductase system that is independent of H_2_ (**Fig. 2**) [6]. *M. barkeri* uses three [NiFe] hydrogenases to mediate H_2_ cycling during methylotrophic methanogenesis (**Fig. 1**) [4]. Cytosolic Frh (<u>F</u>_420_-<u>r</u>educing <u>h</u>ydrogenase), encoded by the *frhADGB* operon, functions to oxidize reduced coenzyme F_420_ (F_420_H_2_), produced by the first two methyl oxidation steps, to form H_2_ inside the cell. The final methyl oxidation step results in the reduction of the electron carrier protein ferredoxin. Reduced ferredoxin is oxidized by the membrane-bound <u>e</u>nergy-<u>c</u>onverting <u>h</u>ydrogenase (Ech) that pumps protons outside the cell while producing intracellular H_2_. The H_2_ produced by Frh and Ech is oxidized on the outside of the cell by <u>v</u>iologen-<u>r</u>educing hydrogenase two (Vht). Vht reduces methanophenazine with the electrons derived from H_2_. Finally, methanophenazine provides electrons to heterodisulfide reductase, which in turn reduces CoMS-SCoB. The net result is the generation of a proton gradient that drives ATP production by ATP synthase [4].

**Figure 1.**
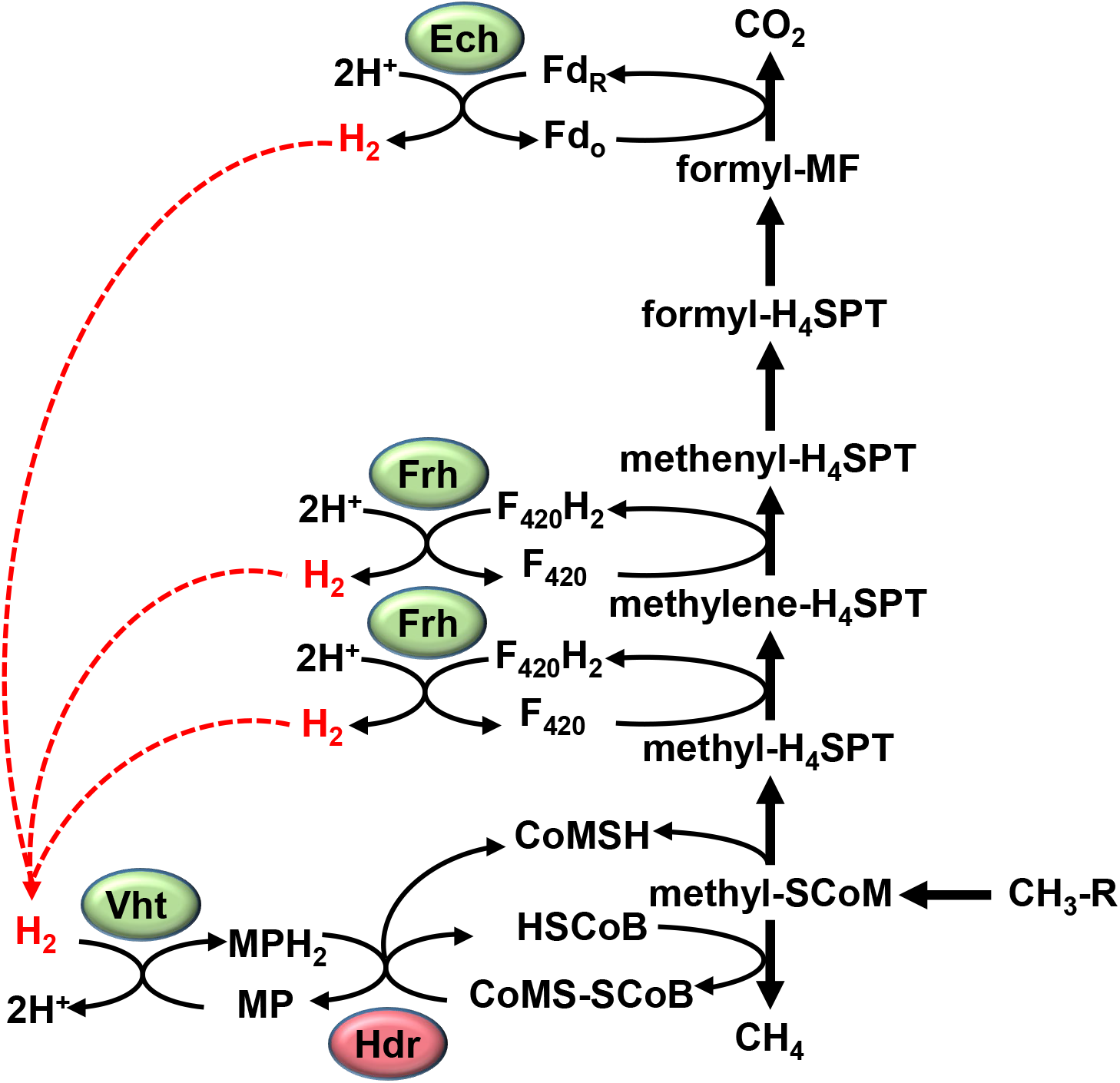
The methylotrophic pathway in *M. barkeri* involves H_2_ cycling. Only enzymes relevant to H_2_ cycling are shown. One methyl group is oxidized to CO_2_ providing the electrons for the indirect reduction of three other methyl groups to CH_4_. Frh hydrogenase produces cytosolic H_2_ by the oxidation of F_420_H_2_ generated during the oxidation of the methyl group. Ech hydrogenase also produces cytosolic H_2_ by the oxidation of Fd_R_. Vht hydrogenase oxidizes H_2_ on the outside of the cell and transfers electrons to MP in the membrane, which then transfers electrons to heterodisulfide reductase (Hdr) to reduce CoMS-SCoB. Fd_o_, oxidized ferredoxin; Fd_r_, reduced ferredoxin; F_420_, coenzyme F_420_; H_4_SPT, tetrahydrosarcinapterin; CoMSH, coenzyme M; HSCoB, coenzyme B; MP, methanophenazine; Hdr, heterodisulfide reductase.

**Figure 2.**
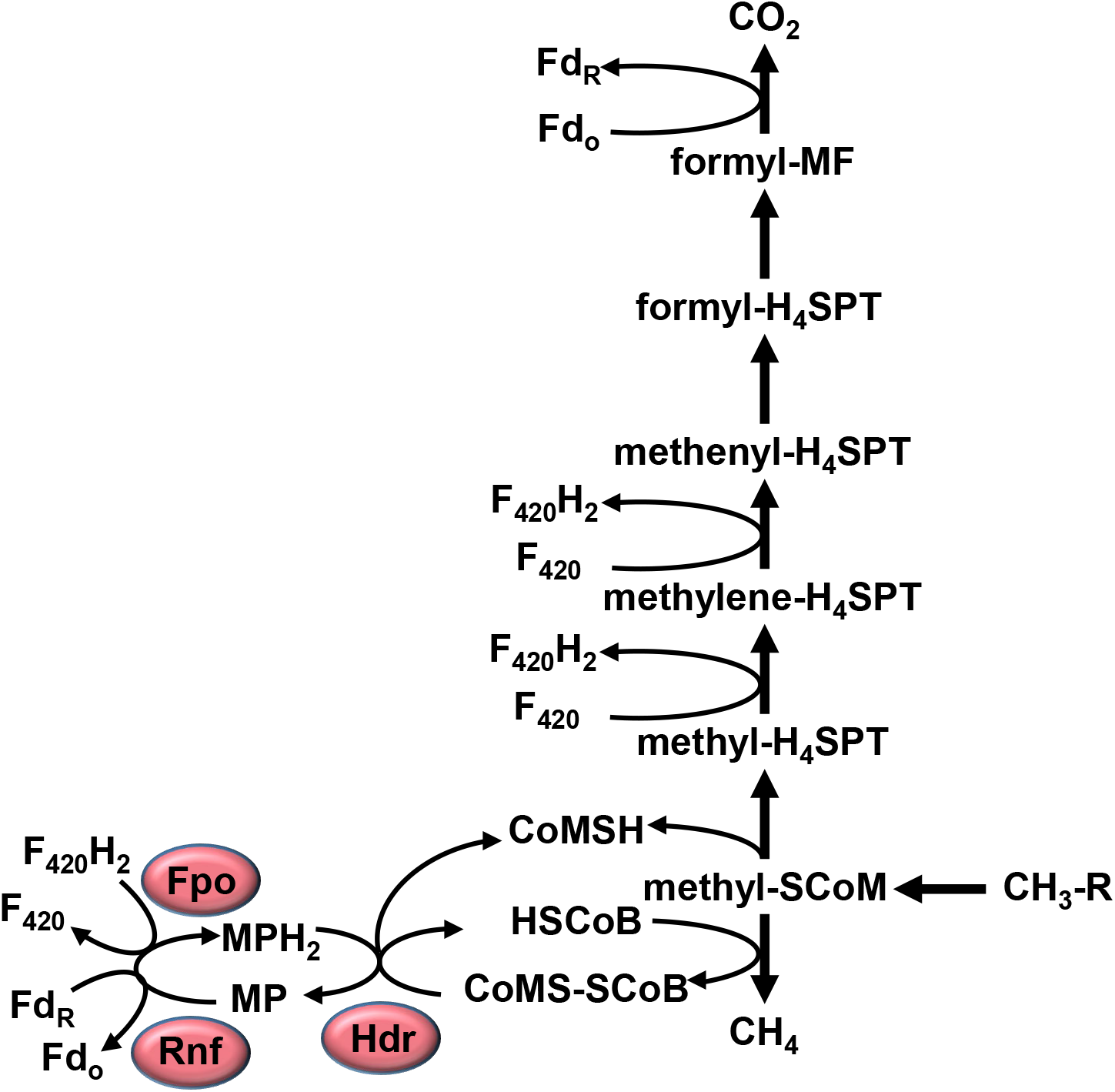
The methylotrophic pathway in *M. acetivorans* does not involve H_2_ cycling. One methyl group is oxidized to CO_2_ providing the electrons for the indirect reduction of three other methyl groups to CH_4_. Fpo oxidizes F_420_H_2_ and Rnf oxidizes Fd_R_ and each transfers electrons to MP in the membrane, which then transfers electrons to heterodisulfide reductase (Hdr) to reduce CoMS-SCoB. Fd_o_, oxidized ferredoxin; Fd_r_, reduced ferredoxin; F_420_, coenzyme F_420_; H_4_SPT, tetrahydrosarcinapterin; CoMSH, coenzyme M; HSCoB, coenzyme B; Fpo, F_420_ dehydrogenase; Rnf, Fd oxidoreductase; MP, oxidized methanophenazine; MPH_2_, reduced methanophenazine Hdr, heterodisulfide reductase.

In contrast, H_2_ cycling is not involved in electron transport chain of *M. acetivorans*. Instead, *M. acetivorans* uses F_420_ dehydrogenase (Fpo) to directly oxidize F_420_H_2_, and the Rnf complex to directly oxidize ferredoxin, with both enzyme complexes reducing methanophenazine (**Fig. 2**) [2, 7]. It is hypothesized that *M. acetivorans* metabolism evolved to be independent of H_2_ due to selective pressure in the marine environment as a result of high levels of sulfate that put methanogens at a competitive disadvantage with sulfate-reducing anaerobes to use H_2_ as an electron donor [8, 9]. Methylotrophic, acetotrophic, and carboxydotrophic growth by *M. acetivorans* does not involve H_2_ as an intermediate [3, 7-10]. However, growth of *M. barkeri* with all substrates involves H_2_ as the preferred intermediate electron carrier. For example, *M. barkeri* contains Fpo that can support methylotrophic growth by means of a F_420_H_2_:CoMS-SCoB oxidoreductase system like *M. acetivorans*, but mutational analyses indicates that the H_2_:CoMS-SCoB oxidoreductase system is the preferred system [11].

Despite methanogenesis by *M. acetivorans* being independent of H_2_, the genome encodes Frh and two copies of the viologen-reducing hydrogenase (Vht and Vhx) [12]. *M. acetivorans* lacks Ech; however, expression of *M. barkeri* Ech in *M. acetivorans* allows *M. acetivorans* to grow by H_2_-dependent methyl reduction, indicating *M. acetivorans* can produce functional hydrogenase [13]. The amino acid sequences of *M. acetivorans* Frh, Vht, and Vhx proteins contain the necessary motifs required for hydrogenase activity. Moreover, *M. acetivorans* contains a *hyp* operon that encodes hydrogenase maturation proteins. However, lysates from wild-type *M. acetivorans* cells exhibit little to no hydrogenase activity under all growth conditions tested [13, 14]. Gene reporter analysis indicates very low transcription of *frh, vht*, and *vhx* operons in *M. acetivorans* [15]. Thus, the importance and role of hydrogenase to *M. acetivorans* has remained a mystery.

The genome of *M. acetivorans* encodes molybdenum (Mo), vanadium (V), and Fe-only (Fe) nitrogenases, the enzymes required for growth using dinitrogen (N_2_) as a nitrogen source (i.e., N_2_ fixation or diazotrophy) [12, 16, 17]. We have recently shown that *M. acetivorans* can grow by fixing N_2_ using Mo-nitrogenase [18]. N_2_ fixation is an energy intensive process; it is estimated that two ATP are hydrolyzed for every electron used by nitrogenase. For example, bacterial Mo-nitrogenase catalyzes the following reaction: N_2_ + 8*e*^*-*^ + 8H^+^ + 16ATP → 2NH_3_ + H_2_ + 16ADP + 16P_i_ [19]. Due to the obligate requirement for the reduction of protons to H_2_ at the expense of electrons and ATP during the reduction of N_2_ by nitrogenase, many diazotrophic bacteria produce an uptake hydrogenase to oxidize H_2_ to recoup energy [20]. To our knowledge, H_2_ production during N_2_ reduction by Mo-nitrogenase from methanogens has not been documented, but methanogen Mo-nitrogenase is highly similar to Mo-nitrogenase from bacteria, and likely also produces H_2_ [21]. Therefore, we hypothesized that *M. acetivorans* utilizes Frh and/or Vht/Vhx to recover energy lost to H_2_ production during N_2_ reduction. To test this hypothesis, we examined the expression and activity of hydrogenase in response to fixed nitrogen availability and used the recently developed CRISPRi-dCas9 system [18] to repress the transcription of the *frh, vht, vhx*, and *hyp* operons in *M. acetivorans* to determine the effect of the loss of hydrogenase expression on diazotrophy and H_2_ metabolism.

## RESULTS

### *M. acetivorans* hydrogenase expression and activity increases during nitrogen fixation

Previous analysis of *M. acetivorans frh, vht, vhx*, and *hyp* operon expression using single-copy reporter gene fusions to each promoter revealed little to no expression of each operon in both *M. acetivorans* and *M. barkeri* cells grown with methanol, methanol + H_2_, or acetate [15]. Therefore, hydrogenase expression in *M. acetivorans* does not appear responsive to growth substrate or the presence of H_2_. To test the hypothesis that expression of hydrogenase(s) and the hydrogenase maturation machinery increases during N_2_ fixation to allow for H_2_ cycling, the transcript abundance of *vhtG, vhxG, frhA*, and *hypD* in *M. acetivorans* cells grown with or without fixed nitrogen (NH_4_Cl) was determined by qPCR. The transcript abundance of all four genes is approximately 2- to 3-fold higher in cells grown with N_2_ compared to cells grown with NH_4_Cl (**Fig. 3**), with only the fold change for *vhtG* not significant. Hydrogenase activity was also compared in lysates from cells grown with or without NH_4_Cl as measured by H_2_-dependent methyl viologen reduction. Lysates from N_2_-grown cells exhibit an approximate 15-fold increase in hydrogenase activity compared to NH_4_Cl-grown cells (**Table 1**). The hydrogenase activity in lysate from N_2_-grown *M. acetivorans* cells is still roughly 6-fold lower than the activity in lysate from NH_4_Cl-grown *M. barkeri* cells (**Table 1**). Overall, these results reveal that one or more of the hydrogenases is functional in *M. acetivorans* and that hydrogenase expression is responsive to fixed nitrogen availability.

**Figure 3.**
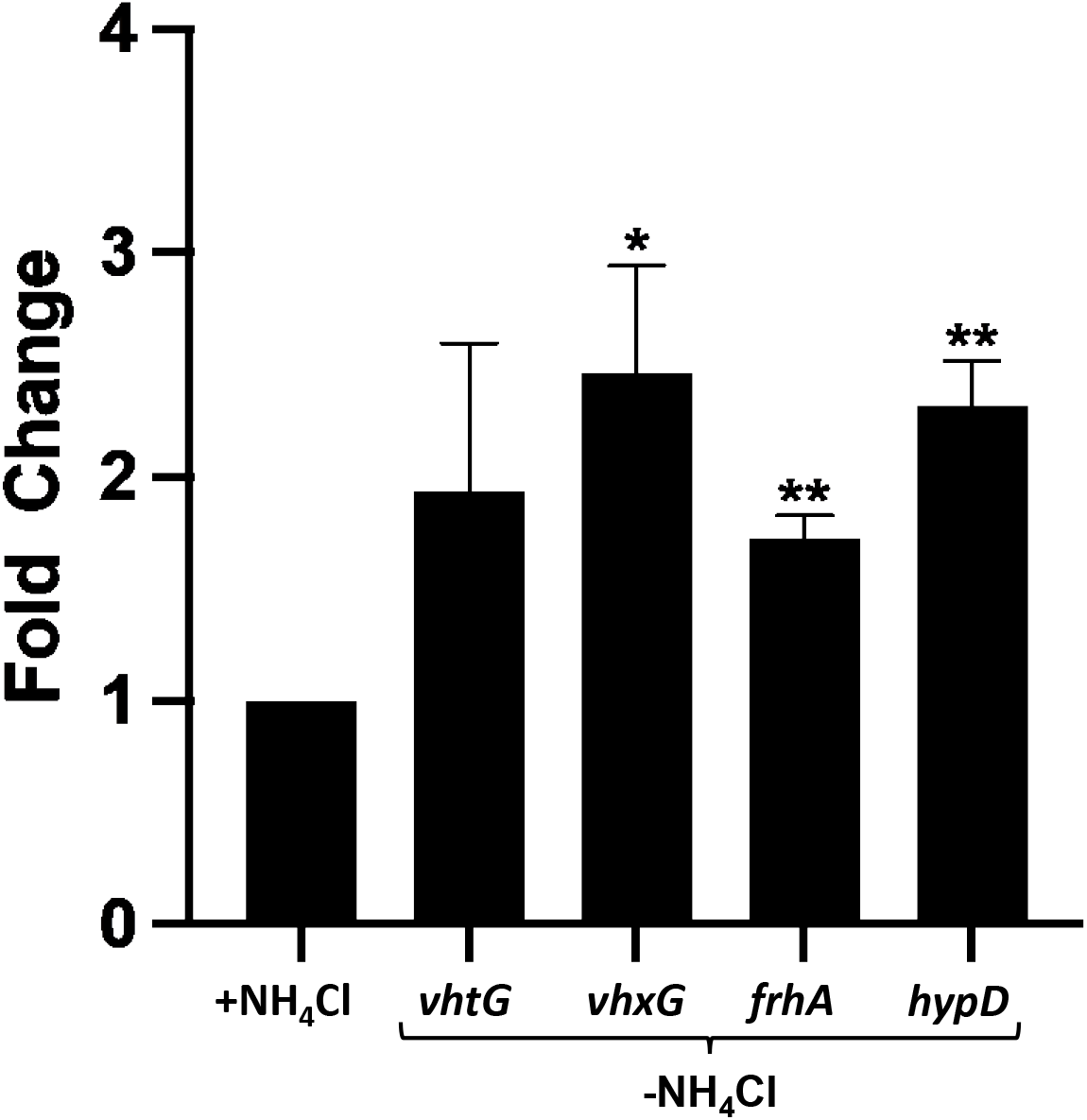
The fold change in *vhtG, vhxG, frhA* and *hypD* transcript abundance in *M. acetivorans* cells grown without NH_4_Cl compared to cells grown with NH_4_Cl. Strain DJL72 was grown in HS medium with 125 mM methanol and with or without NH_4_Cl. qPCR was performed as described in Materials and Methods. The relative abundance of *vhtG, vhxG, frhA, and hypD* in cells grown with NH_4_Cl was set to 1 to determine fold change transcript abundance in cells grown without NH_4_Cl. Data are the means from three biological replicates analyzed in duplicate. *P <0.05, **P <0.01.

**Table 1.** Comparison of methyl-viologen dependent hydrogenase activity in methanogen cell lysates.

| Methanogen | Nitrogen source | Hydrogenase activity<sup>a</sup> |
| --- | --- | --- |
| *M. barkeri* | NH<sub>4</sub>Cl | 1073 ± 38 |
| *M. acetivorans* | NH<sub>4</sub>Cl | 11 ± 3 |
| *M. acetivorans* | N<sub>2</sub> | 172 ± 13 |

<sup>a</sup>H<sub>2</sub>-dependent methyl viologen reduction (nmol×min<sup>-1</sup>×mg<sup>-1</sup>protein). Average of triplicate reactions.

### Expression of Vht hydrogenase is required for H_2_ cycling during nitrogen fixation

To test the importance of each hydrogenase to diazotrophic growth by *M. acetivorans*, the transcription of the *vht, vhx*, and *frh* operons was targeted for repression by dCas9 using the recently developed CRISPRi-dCas9 system [18]. A gRNA was designed to target the coding strand just after the start codon of the first gene in each operon (**Fig. 4**). Each plasmid containing a gRNA and dCas9 was separately integrated into the chromosome of *M. acetivorans* strain WWM73 (pseudo-wild type) to generate strains DJL150-152 (**Fig. 4**). The growth of strains DJL150-152 with or without NH_4_Cl was compared to strain DJL72 (gRNA-free control) with methanol as the carbon and energy source (**Fig. 5**). Strains DJL150-152 grow like control strain DJL72 with NH_4_Cl as the nitrogen source, with no difference in growth rate or final cell density (**Fig. 5A**). Growth of strains DJL151 and DJL152 without NH_4_Cl is also identical to control strain DJL72 (**Fig. 5B**). However, strain DJL150 harboring the gRNA targeting the *vht* operon grows slower and achieves a significantly lower final cell yield than the control (**Fig. 5B**). These results indicate production of Vht, but not Vhx or Frh, is required for optimal diazotrophic growth. The repression of the dCas9-targeted operon in each strain was confirmed by determination of the transcript abundance for *vhtG, vhxG*, and *frhA* by qPCR. Importantly, a greater than 90% reduction in transcript abundance is observed for each targeted gene in cells grown with or without NH_4_Cl (**Fig. 6**). Thus, the lack of an observable phenotype with strains DJL151 and DJL152 is not due to insufficient dCas9-mediated repression of the *vhx* and *frh* operons, respectively.

**Figure 4.**
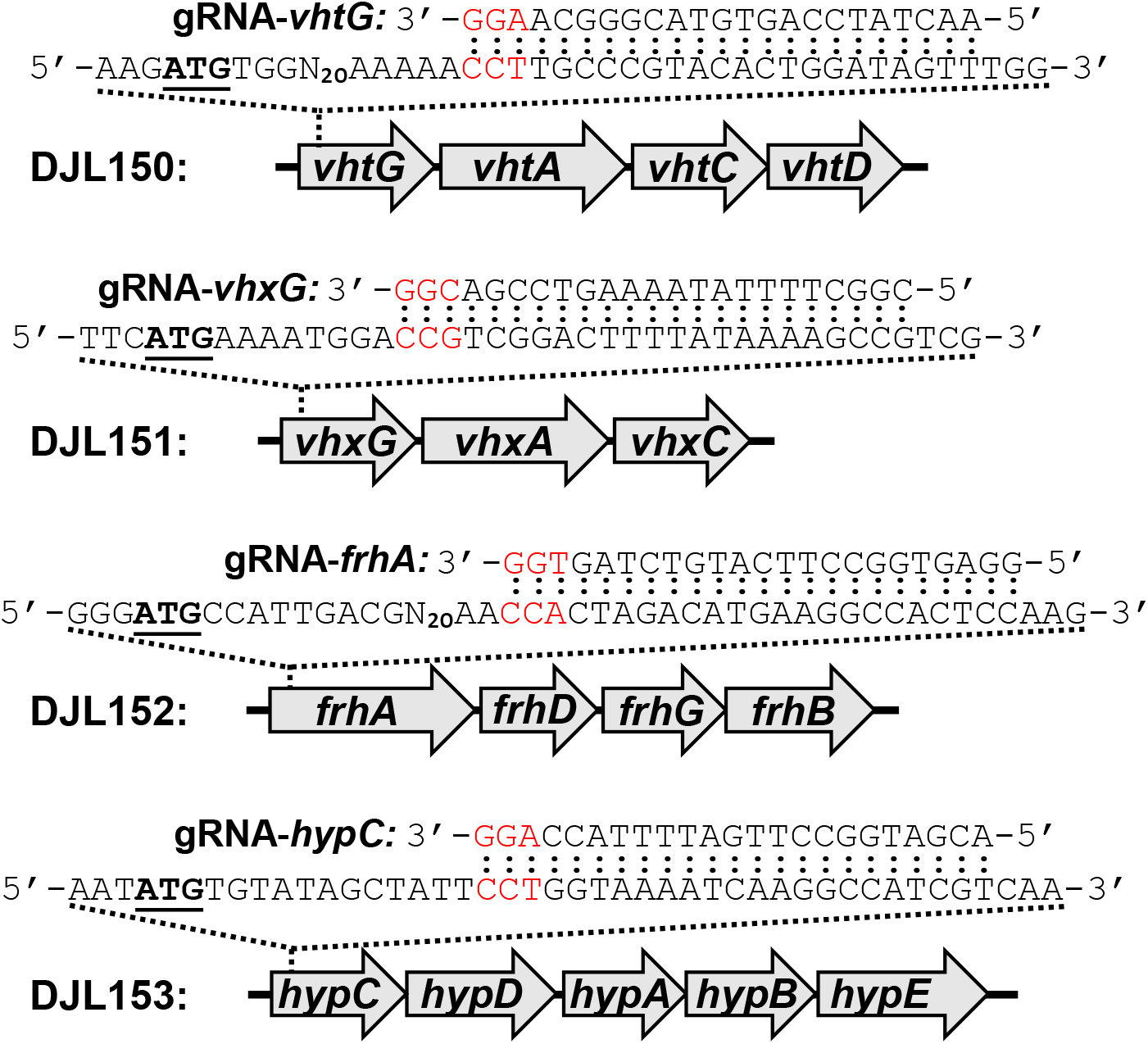
Location of gRNAs targeting the *vht, vhx, frh*, and *hyp* operons for dCas9-mediated repression in *M. acetivorans* strains DJL150, DJL151, DJL152, and DJL153. Start codons are bold and underlined. PAM sequences are red.

**Figure 5.**
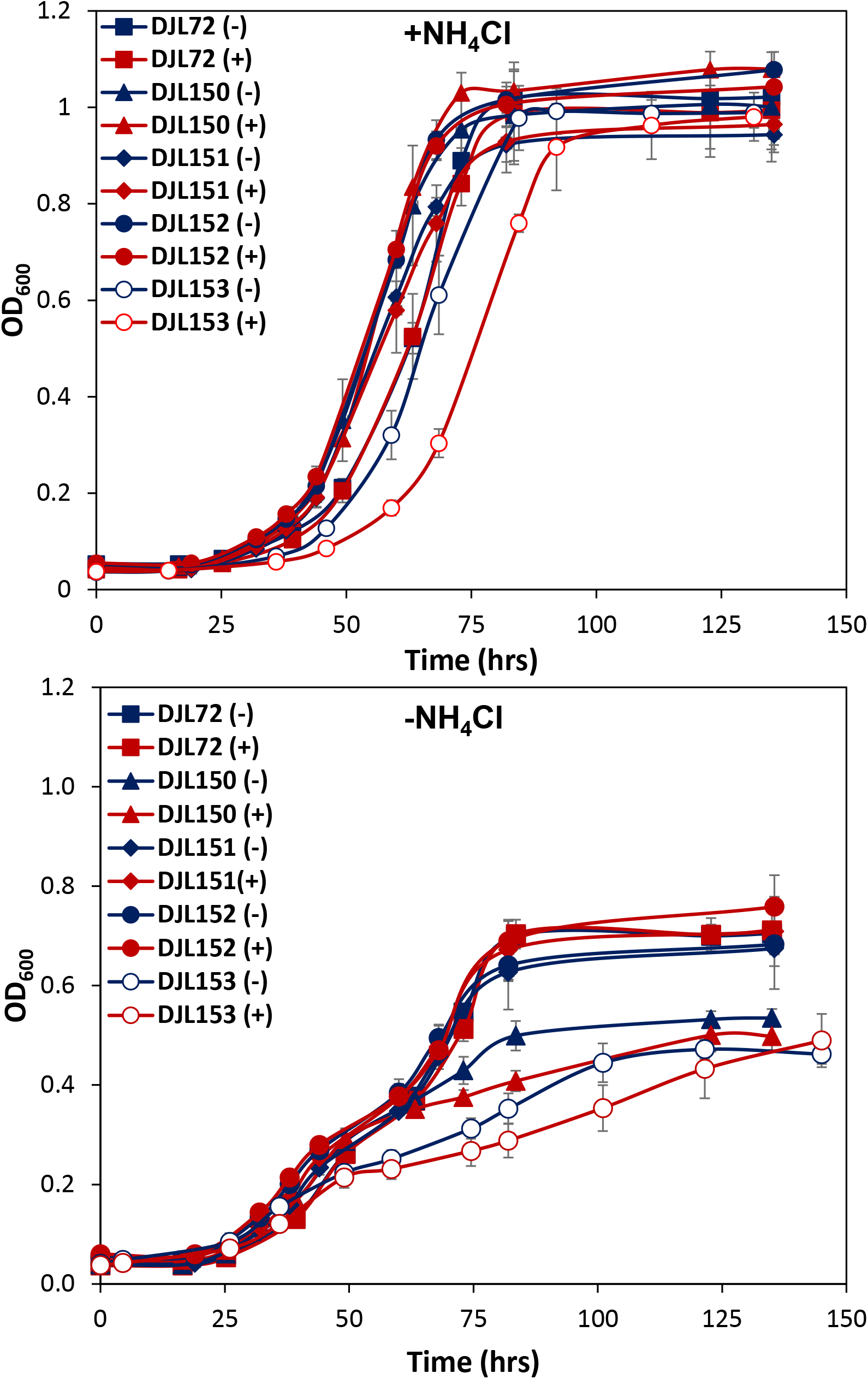
Comparison of the growth of *M. acetivorans* strains DJL150, DJL151, DJL152, and DJL153 to the control strain DJL72 (gRNA-free) with NH_4_Cl or without NH_4_Cl. H_2_ (83 µmoles) was omitted (-) and or added (+) to the headspace of the cultures prior to inoculation.

**Figure 6.**
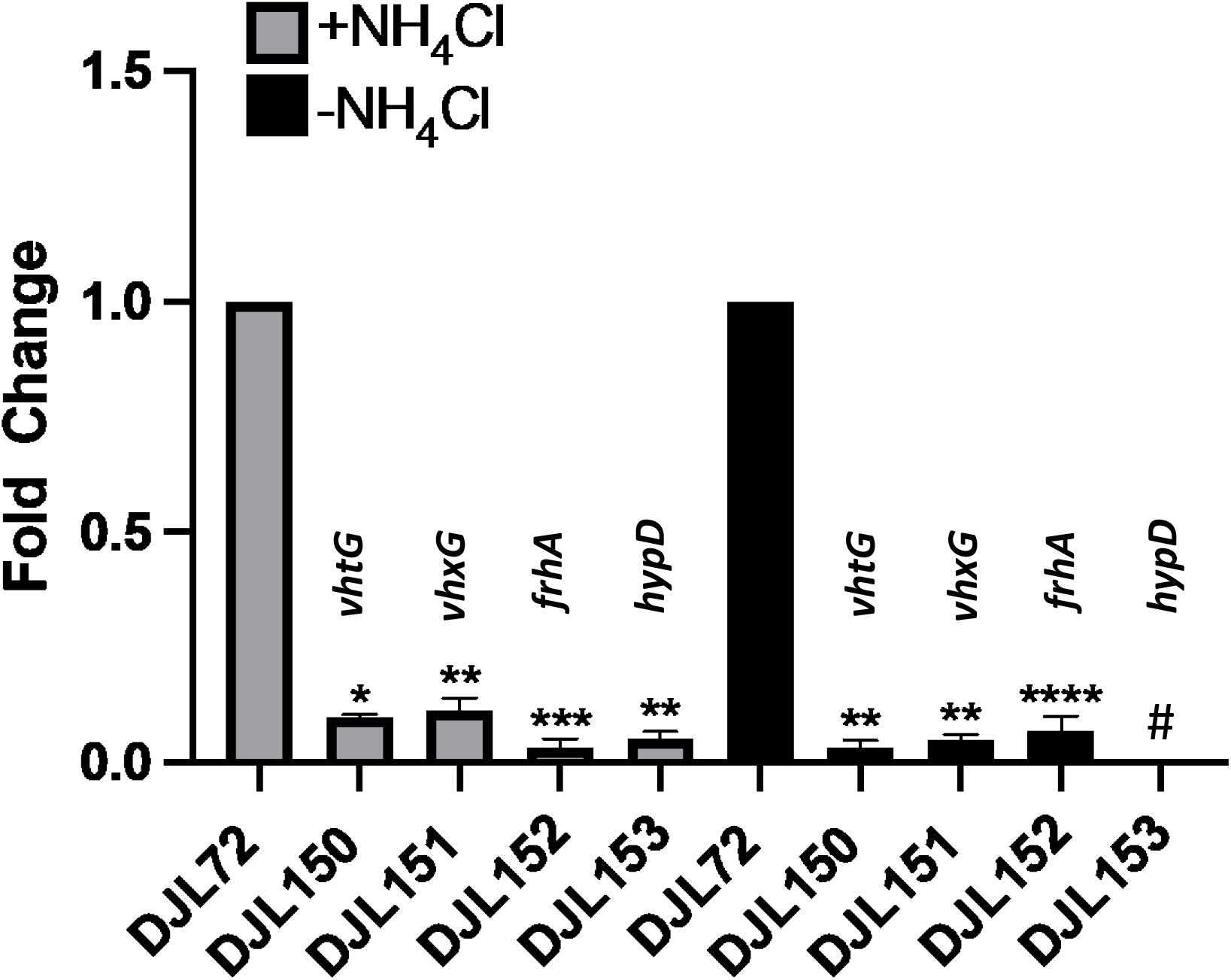
The fold change in relative transcript abundance of genes targeted for dCas9-mediated repression in *M. acetivorans*. Strains were grown in HS medium with 125 mM methanol and with or without NH_4_Cl. qPCR was performed as described in Materials and Methods. Data are the means from three biological replicates analyzed in duplicate. The relative abundance of *vhtG, vhxG, frhA, and hypD* in strain DJL72 (gRNA-less) was set to 1 to determine fold change in the other strains. *, *P* < 0.05; **, *P* < 0.01; ***, *P* < 0.001; ****, *P* < 0.0001. ^#^Transcript abundance of *hypD* was below the detection limit in strain DJL153 grown without NH_4_Cl.

Although H_2_ is produced by nitrogenase during the reduction of N_2_, it is a competitive inhibitor of N_2_ reduction by bacterial Mo-nitrogenase [22-24]. Thus, in addition to recovering energy lost to the production of H_2,_ hydrogenase may also serve to protect nitrogenase from H_2_ inhibition by lowering the level of H_2_. The effect of elevated H_2_ levels on diazotrophic growth of the hydrogenase repression strains was tested by the addition of H_2_ to the headspace of culture tubes prior to inoculation. The addition of exogenous H_2_ does not alter the growth profile of any of the hydrogenase repression strains with NH_4_Cl as the nitrogen source (**Fig. 5A**). The diazotrophic growth (-NH_4_Cl) of strains DJL72, DJL151, and DJL152 with H_2_ in the headspace were identical to the growth profiles of same cultures with no added H_2_ (**Fig. 5B**). However, diazotrophic growth of strain DJL150 with H_2_ in the headspace was slower compared to diazotrophic growth in tubes without additional H_2_, indicating Vht may also protect Mo-nitrogenase from H_2_ inhibition during N_2_ reduction.

Repression of the *vht* operon clearly impairs diazotrophic growth of *M. acetivorans*, which is further hindered by the presence of elevated H_2_ (**Fig. 5**). To determine if loss of *vht* operon expression affects H_2_ metabolism during N_2_ fixation by *M. acetivorans*, the total amount of H_2_ was measured after the cessation of growth in culture tubes with or without added NH_4_Cl and H_2_ (**Table 2**). Control strain DJL72 grown in the absence of additional H_2_ and without NH_4_Cl does not produce a detectable amount of H_2_, indicating that any H_2_ produced endogenously by nitrogenase is subsequently consumed. However, a similar amount of H_2_ is detected in cultures of strain DJL72 grown with or without NH_4_Cl that had H_2_ added prior to inoculation, revealing that *M. acetivorans* does not consume exogenously available H_2_ during non-diazotrophic or diazotrophic growth. Consistent with the growth profiles that matched control strain DJL72 (**Fig. 5**), the amount of H_2_ detected in cultures of strains DJL151 and DJL152 is like that detected in cultures of strain DJL72, revealing loss of *vhx* and *frh* expression does not alter H_2_ metabolism. However, repression of the *vht* operon in strain DJL150 prevents H_2_ consumption. In strain DJL150 culture tubes without added H_2_, there is a detectable amount of H_2_ after diazotrophic growth, whereas no H_2_ is detected after non-diazotrophic growth. Similar levels of H_2_ are detected in non-diazotrophic and diazotrophic cultures of strain DJL150 with added H_2_. These results clearly demonstrate that Mo-nitrogenase produces H_2_ during the reduction of N_2_ within *M. acetivorans* and that Vht hydrogenase, but not Vhx or Frh hydrogenase, is required for the consumption of the produced H_2_. The results also reveal that *M. acetivorans* does not significantly consume exogenously provided H_2_ even when Vht hydrogenase is actively consuming H_2_ produced by nitrogenase.

**Table 2.** H_2_ determination after growth of *M. acetivorans* strains with or without NH_4_Cl in culture tubes with or without initial H_2_.

| *M. acetivorans* strain | Initial H<sub>2</sub> added<sup>a</sup> | Nitrogen source | Final [H<sub>2</sub>]<sup>b</sup> (μmoles) |
| --- | --- | --- | --- |
| DJL72<br>(gRNA-less) | - | NH<sub>4</sub>Cl | 0 |
|  | - | N<sub>2</sub> | 0 |
|  | + | NH<sub>4</sub>Cl | 68 ± 16 |
|  | + | N<sub>2</sub> | 85 ± 10 |
| DJL150<br>(gRNA-*vhtG*) | - | NH<sub>4</sub>Cl | 0 |
|  | - | N<sub>2</sub> | 73 ± 3 |
|  | + | NH<sub>4</sub>Cl | 97 ± 23 |
|  | + | N<sub>2</sub> | 111 ± 29 |
| DJL151<br>(gRNA-*vhxG*) | - | NH<sub>4</sub>Cl | 0 |
|  | - | N<sub>2</sub> | 0 |
|  | + | NH<sub>4</sub>Cl | 73 ± 12 |
|  | + | N<sub>2</sub> | 62 ± 17 |
| DJL152<br>(gRNA-*frhA*) | - | NH<sub>4</sub>Cl | 0 |
|  | - | N<sub>2</sub> | 0 |
|  | + | NH<sub>4</sub>Cl | 72 ± 6 |
|  | + | N<sub>2</sub> | 61 ± 9 |
| DJL153<br>(gRNA-*hypC*) | - | NH<sub>4</sub>Cl | 0 |
|  | - | N<sub>2</sub> | 33 ± 7 |
|  | + | NH<sub>4</sub>Cl | 63 ± 11 |
|  | + | N<sub>2</sub> | 64 ± 19 |

<sup>a</sup>H<sub>2</sub> (83 μmoles) was omitted (-) or added (+) to the headspace of H<sub>2</sub>-free culture tubes as indicated prior to inoculation.
<sup>b</sup>H<sub>2</sub> was determined in headspace of at least triplicate cultures by GC as described in Materials and Methods.

### Expression of the *hyp* operon is required for H_2_ cycling during nitrogen fixation

HypABCDE are essential for the post-translational modification and maturation of [NiFe]-hydrogenases in bacteria [25]. To our knowledge, the importance of Hyp proteins to the maturation of [NiFe]-hydrogenase in methanogens has not been determined. To test the importance of HypABCDE to hydrogenase function in *M. acetivorans*, the *hyp* operon was targeted for dCas9-mediated repression with a gRNA that targets within the coding sequence of *hypC* (**Fig. 4**). Repression of the *hyp* operon in strain DJL153 was confirmed by qPCR analysis of the transcript abundance of *hypD* (**Fig. 6**). Importantly, non-diazotrophic and diazotrophic growth of strain DJL153 is similar to the growth of strain DJL150, with both exhibiting impaired growth with N_2_ but not with NH_4_Cl (**Fig. 5**). The addition of initial H_2_ to the headspace of culture tubes further slows the growth of strain DJL153 with N_2_, like that seen with strain DJL150 (**Fig. 5B**). Moreover, H_2_ is detected in culture tubes of strain DJL153 grown with N_2_ but H_2_ is not detected in culture tubes grown with NH_4_Cl (**Table 2**). These results are consistent with HypABCDE being essential for the maturation of Vht hydrogenase that is required for consumption of H_2_ produced by Mo-nitrogenase.

### Correlation of nitrogenase and hydrogenases in Methanosarcinales

Both *M. barkeri* and *M. mazei* contain Ech and carry out H_2_-dependent methanogenesis, whereas *M. acetivorans* lacks Ech and carries out H_2_-independent methanogenesis, indicating that in the Methanosarcinales the presence or absence of Ech is linked to H_2_-dependent and H_2_-independent methanogenesis, respectively [4]. Therefore, members of the Methanonsarcinales that lack Ech are likely non-hydrogenotrophic but may contain hydrogenase to consume H_2_ produced endogenously by nitrogenase as shown here for *M. acetivorans*. To determine if there is a correlation between the presence of nitrogenase with the presence of Vht and/or Frh in Ech-lacking Methanosarcinales, we surveyed available complete genomes of Methanosarcinales in the NCBI database for the presence of the *nif* (Mo-nitrogenase), *vht, frh*, and *hyp* operons, with results shown in **Table 3**. Of the 20 included species, 10 contain *nif* and are predicted to be capable of diazotrophy. Of the 10 *nif*-containing species, seven species contain *vht*, with four of these species also containing *frh*. There are no species that only contain *frh*. Three species, *Methanolobus psychrophilus* R15, *Methanolobus zinderi*, and *Methanothrix soehngenii* GP6, contain *nif*, but lack both *vht* and *frh*. However, to our knowledge none of the three species have been experimentally shown to fix N_2_, so it is possible the *nif* operon does not encode functional nitrogenase in these methanogens.

**Table 3.**
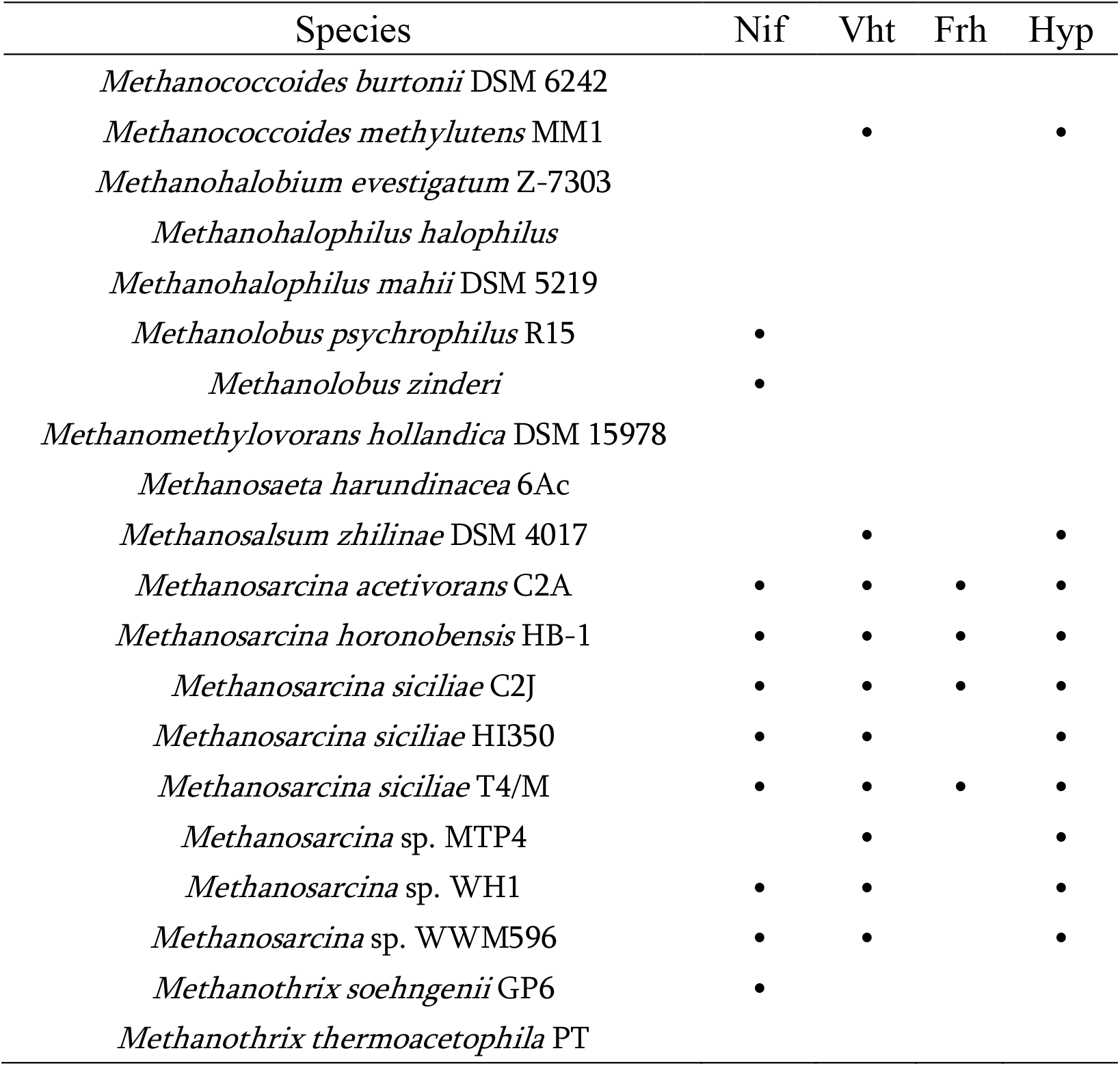
The prevalence of nitrogenase (Nif), hydrogenases (Vht and Frh), and hydrogenase maturation machinery (Hyp) in sequenced Methanosarcinales that lack Ech hydrogenase.

| Species | Nif | Vht | Frh | Hyp |
| --- | --- | --- | --- | --- |
| *Methanococcoides burtonii* DSM 6242 |  |  |  |  |
| *Methanococcoides methylutens* MM1 |  | • |  | • |
| *Methanohalobium evestigatum* Z-7303 |  |  |  |  |
| *Methanohalophilus halophilus* |  |  |  |  |
| *Methanohalophilus mahii* DSM 5219 |  |  |  |  |
| *Methanolobus psychrophilus* R15 | • |  |  |  |
| *Methanolobus zinderi* | • |  |  |  |
| *Methanomethylovorans hollandica* DSM 15978 |  |  |  |  |
| *Methanosaeta harundinacea* 6Ac |  |  |  |  |
| *Methanosalsum zhilinae* DSM 4017 |  | • |  | • |
| *Methanosarcina acetivorans* C2A | • | • | • | • |
| *Methanosarcina horonobensis* HB-1 | • | • | • | • |
| *Methanosarcina siciliae* C2J | • | • | • | • |
| *Methanosarcina siciliae* HI350 | • | • |  | • |
| *Methanosarcina siciliae* T4/M | • | • | • | • |
| *Methanosarcina* sp. MTP4 |  | • |  | • |
| *Methanosarcina* sp. WH1 | • | • |  | • |
| *Methanosarcina* sp. WWM596 | • | • |  | • |
| *Methanotherix soehngeni* GP6 | • |  |  |  |
| *Methanotherix thermoacetophila* PT |  |  |  |  |

Only two species, *Methanococcoides methylutens* MM1 and *Methanosarcina* sp. MTP4, contain *vht* but not *nif*. Importantly, all species that contain *vht* also contain *hyp* that was experimentally shown here to be required for Vht function in *M. acetivorans*. Thus, Vht is likely functional in both *M. methylutens* MM1 and *Methanosarcina* sp. MTP4 and may function in H_2_ cycling independent of nitrogenase. Overall, the results show a correlation between Mo-nitrogenase and Vht hydrogenase in the Methanosarcinales that carry out H_2_-independent methanogenesis, especially within the genus *Methanosarcina* as all species that contain *nif* also contain *vht*.

## DISCUSSION

The results from this study reveal a role for membrane-bound Vht hydrogenase in the consumption of H_2_ produced during the reduction of N_2_ to 2NH_3_ by Mo-nitrogenase in the non-hydrogenotrophic methanogen *M. acetivorans*. It is not surprising that Vht serves as the uptake hydrogenase required for the cycling of H_2_ produced endogenously by nitrogenase. First, H_2_ oxidation is the exclusive function of Vht in *M. barkeri*, whereby Vht oxidizes H_2_ produced by Frh and/or Ech [11]. Both Frh and Ech can either consume or produce H_2_ in *M. barkeri*, depending on growth substrate and environmental conditions [4]. Strains of *M. barkeri* also have nitrogenase and can fix N_2_ [26, 27]; thus, Vht likely also serves to oxidize nitrogenase-produced H_2_ as well as that produced by Ech and/or Frh in *M. barkeri*. Moreover, of the three hydrogenases in *M. barkeri*, only Vht is essential to growth of wild-type cells under all conditions, while Ech and Frh are dispensable under certain growth conditions [11]. Vht is not essential in *M. acetivorans* since no phenotype was observed for *vht* repression strain DJL150 under non-diazotrophic growth conditions. Second, *A vinelandii*, the primary model aerobic diazotrophic bacterium, also contains one membrane bound (Hox) and one cytosolic hydrogenase [28]. Only membrane bound Hox is required for H_2_ consumption during N_2_ fixation by *A. vinelandii* [29, 30]. The role of the cytosolic hydrogenase in *A. vinelandii* is unknown. Deletion of *hox* from *A. vinelandii* results in the detection of nitrogenase-produced H_2_ [31], like that seen here during repression of *vht* in *M. acetivorans*. Finally, it is logical from an energy conservation standpoint that *M. acetivorans* would use Vht to oxidize nitrogenase-produced H_2_, since oxidation of H_2_ by Vht directly contributes to the generation of a proton gradient that can be used for ATP synthesis [4].

Based on current knowledge of the physiology of *Methanosarcina* sp. and the results in this study, we propose the model shown in **Figure 7** for electron transfer and energy conservation during diazotrophic growth with methanol by *M. acetivorans*. The typical methylotrophic pathway is used to oxidize and reduce methyl groups of methanol to CO_2_ and CH_4_ (**Fig. 2**). However, during diazotrophy *M. acetivorans* now employs a bifurcated electron transport chain with H_2_-dependent and H_2_-independent branches like *M. barkeri* [11]. Since ferredoxin and flavodoxin are the known electron donors to nitrogenase in bacteria [16], it is likely that much of the reduced ferredoxin generated from the last methyl oxidation step is used by nitrogenase in addition to other enzymes that carry out reductive biosynthesis. Therefore, Fpo dehydrogenase oxidation of F_420_H_2_ is likely the primary entry point for electrons into the H_2_-independent branch and Rnf oxidation of reduced ferredoxin is minimal or does not occur. ATP-dependent reduction of N_2_ and H^+^ to 2NH_3_ and H_2_ by nitrogenase provides NH_3_ for biosynthesis and the produced H_2_ is oxidized on the outside of the membrane by Vht, which constitutes the H_2_-dependent branch. Both branches contribute to the proton gradient needed for ATP production by ATP synthase. This model is supported by the detection of H_2_ and the lower cell yields in culture tubes containing strains DJL150 or DJL153 growing with N_2_ (**Table 2 and Fig. 5B**). Since there is an obligate requirement for nitrogenase to produce H_2_ during the reduction of N_2_, it is also likely that Vht is required for H_2_-cycling during diazotrophic growth of *M. acetivorans* with acetate, which is likely the predominant substrate in nature [1, 3].

**Figure 7.**
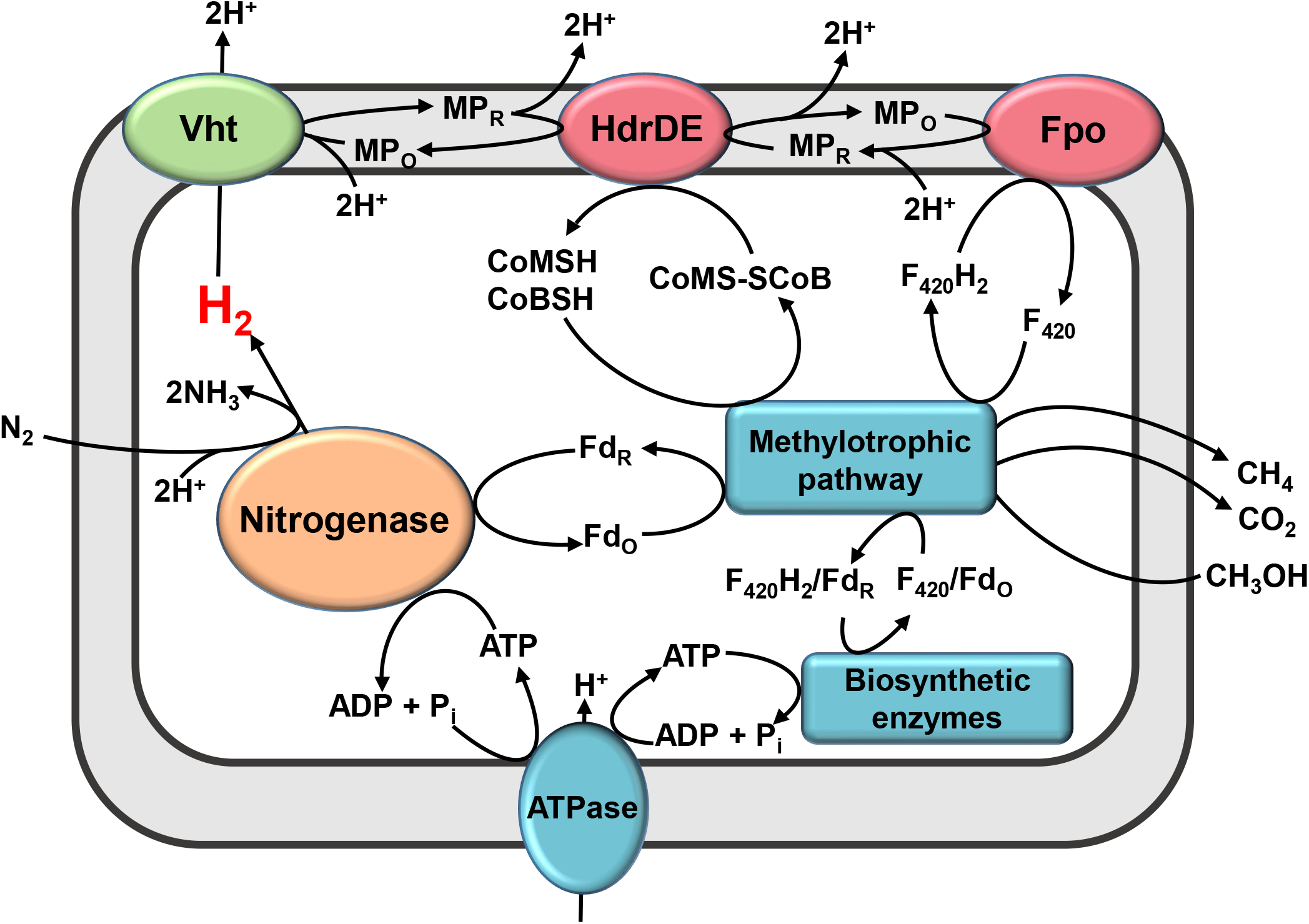
Model of electron transfer and energy conservation during diazotrophy by *M. acetivorans* with methanol. The methylotrophic pathway (as shown in Fig. 2) allows methyl oxidation and reduction to CO_2_ and CH_4_, respectively. Electrons are transferred via ferredoxin to nitrogenase to supported ATP-dependent reduction of N_2_ and 2H^+^. A branched electron transport chain is used to generate a H^+^ gradient. The H_2_-dependent branch requires Vht oxidation of H_2_ and H_2_-independent branch requires Fpo oxidization of F_420_H_2_. Both branches transfer electrons to MP in the membrane, which then transfers electrons to heterodisulfide reductase (Hdr) to reduce CoMS-SCoB. Fd_o_, oxidized ferredoxin; Fd_r_, reduced ferredoxin; F_420_, coenzyme F_420_; HSCoM, coenzyme M; HSCoB, coenzyme B; MP, methanophenazine; Hdr, heterodisulfide reductase.

Vht is clearly required for consumption of endogenously produced H_2_ by nitrogenase in *M. acetivorans*. Vht also likely protects nitrogenase from inhibition by H_2_ during N_2_ reduction by lowering the local concentration of H_2_, since diazotrophic growth of strain DJL150 is depressed in culture tubes with additional H_2_ (**Fig. 5B**). However, the results indicate *M. acetivorans* does not consume additional exogenous H_2_ during diazotrophic growth, despite Vht being active, as evidenced by similar amounts of H_2_ present in both non-diazotrophic and diazotrophic cultures of strains DJL72 (control), DHL151 (gRNA*-vhx*), and DJL152 (gRNA*-frh*) (**Table 2**). Thus, the consumption of H_2_ appears stoichiometric with the amount of N_2_ reduced, despite the potential for the oxidation of exogenous H_2_ contributing to the proton gradient that could increase ATP generation. A possible explanation for the lack of exogenous H_2_ consumption is that since *M. acetivorans* lacks Ech, it does not have a mechanism for H_2_-dependent reduction of ferredoxin that is needed for biosynthesis, unlike *M. barkeri*. During methylotrophic methanogenesis by *M. acetivorans*, the primary and likely only mechanism to reduce ferredoxin is by the last step in the methyl oxidation pathway (**Fig. 2**). Therefore, *M. acetivorans* must continue to completely oxidize methyl groups to generate reduced ferredoxin needed for nitrogenase and biosynthetic reactions (e.g., pyruvate synthesis) (**Fig. 7**). The first two steps require the reduction of F_420_, which must be oxidized for continued methyl oxidation to occur. The primary mechanism to oxidize F_420_H_2_ during methylotrophic methanogenesis by *M. acetivorans* is by Fpo of the F_420_H_2_:CoBS-SCoM electron transport chain. Both Vht and Fpo reduce the same electron acceptor, methanophenazine (**Fig. 7**). Consequently, if more H_2_ was oxidized by Vht this would likely prevent Fpo from oxidizing F_420_H_2_ which would slow or shut down the methyl oxidation pathway that ultimately inhibits diazotrophic growth. How *M. acetivorans* controls and balances electron flux through Vht in response to fixed nitrogen levels and elevated exogenous H_2_ is unclear but warrants further investigation.

The maturation of [NiFe]-hydrogenases has been extensively studied in bacteria (e.g., *E. coli*) and some archaea (e.g., *Thermococcus kodakarensis*) [32, 33]. HypC, HypD, HypE and HypF are required for the assembly and delivery of the iron center and HypA and HypB are required for the insertion of nickel into the active site. [NiFe]-hydrogenases are critical to methanogenesis, yet to our knowledge the requirement of HypABCDE for [NiFe]-hydrogenase maturation has not been experimentally verified in any methanogen. This is likely due to [NiFe]-hydrogenases being essential in virtually all methanogens preventing the ability to delete or mutate *hypABCDE* [34]. Here we show that dCas9-mediated repression of the *hypCDABE* operon in *M. acetivorans* prevents H_2_ cycling, which provides the first experimental evidence that HypABCDE is required for functional hydrogenase in a methanogen. Moreover, since [NiFe]-hydrogenases, including Vht, are not essential to *M. acetivorans* it can serve as a platform to investigate hydrogenase maturation in methanogens.

The results in this study do not provide evidence of a role for Vhx or Frh in the physiology of *M. acetivorans. M. barkeri* also contains Vhx and prior investigation indicates Vhx is not functional in *M. barkeri* [4, 11, 15]. Thus, it is also likely that Vhx does not function in the physiology of *M. acetivorans*. In *M. barkeri*, the activity of Frh is reversible allowing Frh to balance the intracellular levels of H_2_ and F_420_H_2_ depending on growth conditions [4]. During diazotrophic growth by *M. acetivorans*, it is possible that Frh could oxidize H_2_ and reduce F_420_ that could then transfer electrons to Fpo. However, the results indicate this is not occurring since nitrogenase-produced H_2_ is detected in culture tubes containing strain DJL150, where Frh should be active, and strain DJL153, where Frh should be inactive due to repression of *hyp* (**Table 2**). It is still possible that Frh serves to balance the H_2_:F_420_H_2_ ratio in *M. acetivorans* during diazotrophic growth, a process that is likely not essential and therefore does manifest an impaired diazotrophic growth phenotype in strain DJL152. Alternatively, Frh may be needed under conditions with increased H_2_ production. In bacteria, the ATP-dependent production of H_2_ during N_2_ reduction by V-nitrogenase and Fe-nitrogenase is higher than Mo-nitrogenase. Bacterial V-nitrogenase and Fe-nitrogenase are estimated to produce 3H_2_ and 7H_2_ for every N_2_ reduced, respectively [35]. Thus, it is possible Frh becomes important when *M. acetivorans* fixes N_2_ using the alternative nitrogenases, and we are currently testing this hypothesis.

The detection of H_2_ in the headspace of culture tubes containing the *vht* and *hyp* repression strains growing by N_2_ fixation is significant for several reasons. First, to our knowledge, these results are the first to demonstrate the *in vivo* production of H_2_ during the reduction of N_2_ by Mo-nitrogenase in a methanogen, revealing the reaction mechanism of archaeal nitrogenase requires obligate H_2_ production like bacterial nitrogenase [35]. Second, detection of H_2_ as product of *M. acetivorans* nitrogenase activity provides a new method to monitor *in vivo* nitrogenase activity in methanogens. Current methods rely on measuring nitrogenase produced NH_3_, which is notoriously difficult to assay due to poor sensitivity and high background, or by measuring acetylene reduction to ethylene [36]. Although acetylene reduction assays are simple and convenient since the substrate and product are gases, acetylene inhibits methanogenesis making it difficult to measure *in vivo* nitrogenase activity since methanogenesis is required to provide ATP and electrons to support nitrogenase activity [26, 27]. Conversely, H^+^ reduction to H_2_ by nitrogenase has been used to measure nitrogenase activity in bacteria that lack functional hydrogenase [37]. Using H^+^ reduction to H_2_ to determine *in vivo* nitrogenase activity is not possible with most methanogens due to hydrogenase activity being essential. However, hydrogenase is not essential to *M. acetivorans*, allowing H^+^ reduction to H_2_ in Vht-lacking *M. acetivorans* to be used as method to assay methanogen nitrogenase activity. Third, to our knowledge, the *vht* and *hyp* repression strains of *M. acetivorans* are the first methanogen strains capable of nitrogenase-dependent H_2_ production. H_2_ biofuel production using nitrogenase has been engineered in phototrophic and non-phototrophic bacteria, but not in a methanogen [38-41]. An advantage of nitrogenase over hydrogenase to produce H_2_ biofuel is that the reaction catalyzed by nitrogenase is irreversible unlike that catalyzed by hydrogenase, allowing for stable H_2_ production. The Vht and/or Hyp deficient *M. acetivorans* strains described here provide a foundation to potentially engineer nitrogenase-dependent H_2_ production that exploits the unique physiology of methanogens.

Finally, the results here highlight the utility of the CRISPRi-dCas9 system as simple and efficient mechanism to assess gene function in *M. acetivorans*. A single gRNA was used to repress the transcription of each target operon that achieves near gene deletion results as seen for the repression of the *vht* and *hyp* operons, and previously for the *nif* operon that encodes Mo-nitrogenase [18]. Importantly, CRISPRi-dCas9 strains can be generated in as little as two weeks and are highly stable, whereby tested strains can maintain repression of the target gene/operon for at least one year during passage under non-selective conditions (data not shown).

## Acknowledgments

This work was supported in part by DOE Biosciences grant number DE-SC0019226 (DJL) and NSF grant number MCB1817819 (DJL).

## MATERIALS AND METHODS

### *M. acetivorans* strains and growth

*M. acetivorans* strains used are listed in **Table S2**. *M. acetivorans* strain WWM73 served as the parent strain for all experiments [42]. HS medium was prepared as previously described [43], except NH_4_Cl was omitted. Growth experiments were performed in Balch tubes with 10 ml of HS medium. To remove the residual H_2_ from the headspace of each Balch tube due to medium preparation within an anaerobic chamber, the headspace was flushed at 70 kPa with a N_2_/CO_2_ (80:20) mixture using a syringe filtered (0.22 µM) needle and gas escape needle for two minutes. Removal of H_2_ was confirmed by gas chromatography (see below). Methanol, sulfide, puromycin, and NH_4_Cl were added from anaerobic sterile stocks using sterile syringes prior to inoculation. All *M. acetivorans* strains were grown at 35 °C in Balch tubes containing 10 ml HS medium with 125 mM methanol and 0.025% sulfide. Puromycin (2 µg/ml), and NH_4_Cl (18 mM) were added where indicated. Pure H_2_ gas (83 µmoles) was added using a sterile syringe to the headspace of tubes where indicated. Growth was measured by monitoring optical density at 600 nm (OD_600_) using a spectrophotometer.

### Construction of *M. acetivorans* CRISPRi-dCas9 plasmids and strains

All primers (**Table S1**) and synthetic DNA fragments (gBlocks) were designed using Geneious Prime and purchased from Integrated DNA Technologies (IDT). All plasmids (**Table S1**) to target specific genes and operons for dCas9-mediated repression were constructed by introducing a gBlock that contains a specific gRNA into *Asc*I-digested pDL734 using a Gibson Assembly^®^ Ultra Kit (Cat# GA1200-10, Synthetic Genomics, Inc.) as previously described [18]. *M. acetivorans* strain WWM73 was separately transformed with pDL803, pDL802, pDL801, and pDL800 containing the gRNAs targeting *vht, vhx, frh*, and *hyp*, respectively, using a liposome-mediated transformation protocol as described [44]. Transformants were selected by anaerobic growth on agar plates containing 125 mM methanol and 2 µg/ml puromycin. Colonies were screened by PCR and a positive transformant was selected and designated as the corresponding strain. *M. acetivorans* strains were maintained in HS medium with 125 mM methanol and 2 µg/ml puromycin. *M. acetivorans* strain DJL72 harboring pDL730 without a gRNA was generated previously and used as a control for phenotypic comparisons [18].

### Gene expression analysis

*M. acetivorans* cells (4 ml of culture) were harvested at mid-log phase (OD_600_ = 0.2 to 0.4) by centrifugation at 5,000 × *g* inside an anaerobic chamber (Coy Laboratories). The cell pellets were resuspended in 1ml Trizol (Life Technologies, Cat #15596-026) and stored at –80 °C until use. Total RNA was purified from cells using a Direct-zol RNA Miniprep kit (Zymo research, Cat# R2050) according to the manufacturer’s instructions followed by an additional DNase treatment using DNA-free™ DNA Removal Kit (Thermo Fisher Scientific, Cat# AM1906). RNA concentrations were determined using a Thermo Scientific™ NanoDrop 2000. To determine transcript abundance by quantitative real-time PCR assay (qPCR), cDNA was generated with 300ng of total RNA using iScript™ Select cDNA Synthesis Kit (Bio-Rad, Cat# 1708897) with the following conditions (25 °C for 5 min, 42 °C for 30 min, and 85 °C for 5 min). cDNA was stored at −20 °C until use. Each qPCR reaction (10 µl) contained 1x SsoAdvaced universal SYBR Green Supermix (Bio-Rad, Cat#1725271), 30 nM of forward and reverse primers (**Table S3**), and 300-fold dilution cDNA. The qPCR reactions were performed using a CFX96 Real-time PCR detection system (Bio-Rad) for all samples under the following conditions: 95 °C for 1 min followed by 37 cycles of (95 °C for 10 s, 60 °C for 30s) followed by melting curve analysis to ensure the specificity of amplification. qPCR data were analyzed using ΔΔcq method with 16sRNA used as an internal control. To determine the transcript abundance of a specific gene, Δcq values of the gene in the samples were calibrated to the average of its Δcq values in the control group (DJL72 cells) unless indicated otherwise. Duplicate qPCR reactions were conducted for three biological replicates for each strain.

### H_2_ determination

After the cessation of growth, the total volume of gas produced by each culture tube was measured using a glass syringe, which also normalized the pressure to 1 atm. The total amount of H_2_ in each tube was determined by injection of 50 µl of headspace gas into a Shimazdu Nexis GC-2030 gas chromatograph fitted with a Rt-Q-BOND fused silica PLOT column with a 0.32 mm internal diameter, a 30 m length, and a 10.00 µm film thickness (Restek, VWR #89166-308) and BID detector. The sample split ratio was 42.6, and the carrier gas was helium at 4.44 mL/min. The injection port temperature was 100 °C, column temperature 27 °C, and BID temperature 220 °C. Peak integration was performed using Shimadzu LabSolutions software and moles of H_2_ were determined using a curve generated with H_2_ standards.

### Hydrogenase activity assays

*M. acetivorans* strain WWM73 and *M. barkeri* strain WWM84 were grown in 100 ml of HS medium in 250 ml serum bottles to an OD_600_ of 0.5-0.7 and cells harvested by anaerobic centrifugation (16,000 x *g* for 10 minutes). All subsequent steps were carried out in an anaerobic chamber (Coy Labs) with an atmospheric gas composition of 95% N_2_ and 5% H_2_. Cells were resuspended in 1 mL anoxic buffer (50 mM Tris pH 7.2, 150 mM NaCl, 1 mM benzamidine HCl, 1 mM phenylmethylsulfonyl fluoride) and stored in stoppered 2 ml serum vials at −80 °C until use. Thawed *Methanosarcina* cell resuspensions were sonicated on ice with three 10-second pulses followed by centrifugation at 16,000 x *g* for 10 minutes to remove unbroken cells and debris. The supernatant containing cell-free lysate was transferred to a 2 mL serum vial fitted with a rubber septum. To reductively activate hydrogenase, sodium dithionite prepared in 0.013 M sodium hydroxide was added to a final concentration of 1 mM to each cell-free lysate, followed by a 5 min incubation at 25 °C. To remove residual H_2_ from the vials, the stoppered vials were purged with pure N_2_ for 3 mins using a needle fitted to a gas line and a gas escape needle. H_2_-dependent methyl viologen reduction assays were performed in clear anoxic screw-top 1.8 mL autosampler vials fitted with a silicone-PTFE septum. Each vial contained 1 mL of reaction mix comprising 50 mM Tris pH 7.2, 150 mM NaCl, 20 mM methyl viologen, 0.1 mM dithiothreitol. To remove H_2_, each reaction vial was purged with pure N_2_ for 3 mins using a needle fitted to a gas line and a gas escape needle. One mL of pure H_2_ was added to experimental reaction vials using a gas-tight syringe. Similarly, 1 mL of pure N_2_ was added to control (H_2_-free) reaction vials. Reactions were initiated by the addition of 25 µL of cell-free lysate (30-150 µg of total protein). Methyl viologen reduction was monitored at 578 nm using a Cary spectrophotometer with modified cuvette holders to fit the 1.8 ml autosampler vials. H_2_-dependent methyl viologen specific activity (nmol× min^-1^×mg^-1^ protein) was calculated by subtracting the background reduction from H_2_-free control reactions and using an extinction coefficient of 9780 M^-1^ cm^-1^. Protein concentrations were determined using the Bradford method [45].

### Bioinformatics

Blastp from NCBI was used to search the sequenced genomes of all members of the Methanosarcinales for homologs of characterized nitrogenase (Nif) and Ech, Frh, and Vho/t hydrogenases. The query sequences used were EchA from *Methanosarcina barkeri str. Fusaro* (WP_011305192.1), FrhA from *M. barkeri str. Fusaro* (WP_011305486.1), VhoA from *Methanosarcina mazei Go1 (*WP_011034101.1), and NifD from *M. acetivorans* (WP_011023794.1). Hits with an e-value below 10^−20^ were selected for further investigation to assess the presence of complete operons encoding putative enzyme complexes.

### Data availability

Data is available upon request from the corresponding author.

